# Structural impact of synonymous mutations in six SARS-CoV-2 Variants of Concern

**DOI:** 10.1101/2024.07.06.602340

**Authors:** Alison Ziesel, Hosna Jabbari

## Abstract

SARS-CoV-2 continues to spread and infect people worldwide. While most effort into characterizing variants of this virus have focused on non-synonymous changes, accumulation of synonymous mutations in different viral variants has also occurred. Here we characterize six Variants of Concern in terms of their mutational content, and make predictions regarding the impact of those mutations on potential genomic RNA secondary structure and stability. We find that synonymous mutations typically have no or modest impact to RNA secondary structure. As synonymous mutations are free from the selective pressure imposed on protein-altering mutations, the impact of synonymous mutations is largely limited to RNA secondary structure considerations. The absence of major, structure-altering synonymous mutations emphasize the importance of RNA structure, including within coding regions, to viral fitness.

## Introduction

SARS-CoV-2, the causative agent of the COVID-19 pandemic, is a single stranded RNA virus belonging to the clade Betacoronaviridae [72]. Its original host species may have been horseshoe bats, as the most closely related viruses to SARS-CoV-2 infect those species [1, 24]. SARS-CoV-2 is also closely related to SARS-CoV, the causative agent of the SARS outbreak of 2002-2004 and shares a common receptor with SARS-CoV, the angiotensin converting enzyme 2 (ACE2) protein [30, 61, 64].

As the pandemic has proceeded, a number of novel Variants of Concern (VoCs) have arisen in different geographic regions over time. The World Health Organization (WHO) includes the following features as characteristic of a Variant of Concern [66]:

- exhibits genetic changes associated with increased transmissibility, virulence, immune evasion, susceptibility to therapeutics/detectability
- observed to have a growth advantage over other circulating variants in more than one WHO region
- detrimental changes in clinical disease severity, in epidemiology impacting health systems, or decrease in vaccine effectiveness

In the case of SARS-CoV-2, VoCs are typically described in terms of their characteristic protein-altering, i.e. non-synonymous mutations, particularly protein-changing mutations of the Spike protein. Information on non-coding sequences (such as untranslated regions or intergenic regions) or synonymous mutations within coding regions is typically not provided. Here we summarize six recent VoCs in terms of their described mutations and clinical features. An excellent review of the major VoCs prior to the summer of 2021 is available [56]. A comprehensive overview of the characteristic non-synonymous mutations of all VoCs, Variants of Interest (VoIs), and sublineages of SARS-CoV-2 may be viewed at outbreak.info [14]. Figure 1 shows the phylogenetic relationship of six major VoCs, detailed below, with the reference genome sequence for SARS-CoV-2.

**Fig 1.**
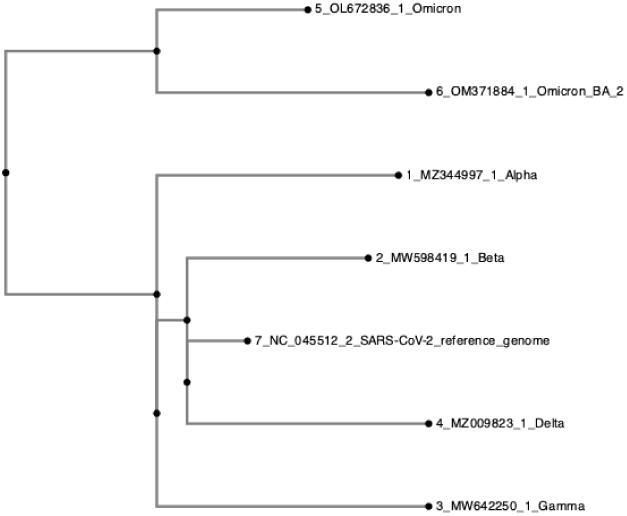
Phylogenetic tree indicating the relationship between six major VoCs and the reference genome for SARS-CoV-2.

### Alpha

The first major VoC, Alpha or B.1.1.7, was first identified in the late summer of 2020 [48]. It is characterized primarily by its mutations N501Y, P618H and del:69-70 of the Spike gene which enhance transmission and infectivity, may enhance cleavage of the Spike surface protein into its S1 and S2 components leading to enhanced viral entry, and may compensate for immune escape mutations respectively [4, 7, 19, 34, 37, 39, 44, 48, 54].

### Beta

The Beta or B.1.351 variant emerged in late 2020 and includes the Spike mutations K417N, E484K and N501Y which each enhance binding of Spike to its receptor ACE2 [28, 58]. Both Alpha and Beta were found to have increased transmissibility relative to the original SARS-CoV-2 strain [58, 63].

### Gamma

The Gamma or P.1 variant was first characterized in late 2020, and includes the Spike mutations N501Y, E484K and K417T; similarly Gamma was observed to be highly transmissible relative to the original strain of SARS-CoV-2 [13].

### Delta

The Delta or B.1.617.2 variant exhibits the characteristic Spike mutations L452R, P681R, T478K and E484Q, which may facilitate antibody evasion, improve viral fusion with the host receptor, may enhance host receptor binding, and act as an escape mutation [11, 53, 55, 65, 67]. Delta exhibits both increased transmissibility as well as reduced susceptibility to monoclonal antibodies approved for emergency use by the American Food and Drug Administration (FDA) [35, 46].

### Omicron

In November 2021, the Omicron or B.1.1.529 variant was designated as a VoC. Omicron and its subvariants are the most highly mutated VoC to date, with at least 32 mutations in the Spike protein including N501Y, N679K and P681H [41, 59, 68].

### Omicron BA.2

The closely related subvariant Omicron BA.2 was first identified in late 2021 and includes a different but overlapping spectrum of mutations with its parent Omicron strain [59]. *In vitro* work with ACE2-expressing 293T cells has demonstrated that Omicron BA.2 is more infectious than Omicron and non-Omicron, D614G-bearing SARS-CoV-2 variants, but is not significantly more prone to immune escape than Omicron or D614G-bearing SARS-CoV-2 variants in a study of monovalent or bivalent mRNA-vaccinated sera [47].

### SARS-CoV-2 genomic organization

As a single stranded RNA virus, SARS-CoV-2 and other coronaviruses are known to form RNA secondary structures. RNA secondary structures are formed when RNA molecules form intra- or intermolecular base pairs, which generates discrete regions of double stranded RNA referred to as stems or helices. Regions of unpaired RNA nucleotides isolated by a stem are referred to as loops. RNA secondary structures with a lower minimum free energy (MFE) are more stable than those structures with a higher MFE, and are more likely to be observed. Studies in SARS-CoV, SARS-CoV-2, and related viruses including murine hepatic virus (MHV), bovine coronavirus (BCov) and others have indicated organized structure in the 5’ and 3’ untranslated regions (UTRs) [6, 17, 20, 22, 23, 29, 31, 33, 38, 51, 60, 70, 71, 73, 74] as well as within the frameshifting stimulatory element (FSE) [5, 15, 27].

The 5’ UTR was characterized largely on work done in MHV and it is predicted to contain five stem-loop (SL) structures, each of which play a role in the viral life cycle. These five stem-loops are believed to be involved in the formation of subgenomic RNAs (sgRNAs) and in the initiation of translation of ORF1ab [20, 29, 31, 33, 38, 71, 74]. SL1 is believed to be involved in a number of key viral functions including protection from Nsp1-mediated translational repression [6]. SL2 is implicated in the production of sgRNAs, and exhibits a conserved YUUGY pentanucleotide loop [29, 33]. SL3 contains the transcription regulatory sequence leader element, and is predicted to unfold to expose that element for sgRNA synthesis [33, 74]. SL4 consists of two stems separated by a bulge, and may act as a spacer element as its deletion is lethal in studies of MHV [73]. Finally, SL5 is a multibranched stem loop including three terminal stem loops named SL5a through SL5c, and includes the start codon for ORF1a, indicating that this structure must be unfolded to begin translation. The exact function of SL5 is not yet understood, although mutations of SL5a can impact viral replication efficiency [20].

Similarly the 3’ UTR exhibits organized structure: shortly after the stop codon of the N gene, the 3’-most gene of the SARS-CoV-2 genome, the 3’ UTR bulged stem-loop structure (BSL) begins, consisting of four stems interspersed with three unpaired bulged subsequences and terminating in a loop [22, 23]. The base of the BSL may alternately interact with an adjacent hairpin structure forming a pseudoknot, where either the pseudoknot or the base of the BSL may be formed, but not both simultaneously; it is thought that this arrangement functions as a molecular toggle switch, oscillating between states to regulate functions pertaining to viral replication [17, 70]. The adjacent hairpin, known as P2, is part of a larger, poorly conserved region known as the hypervariable region (HVR). The 3’-most element of the SARS-CoV-2 genome is the *stem-loop II* -like motif element (s2m), which may play a role in host translation interference [51, 60]. The structures and relative locations of the UTRs and FSE are depicted in Figure 2.

**Fig 2.**
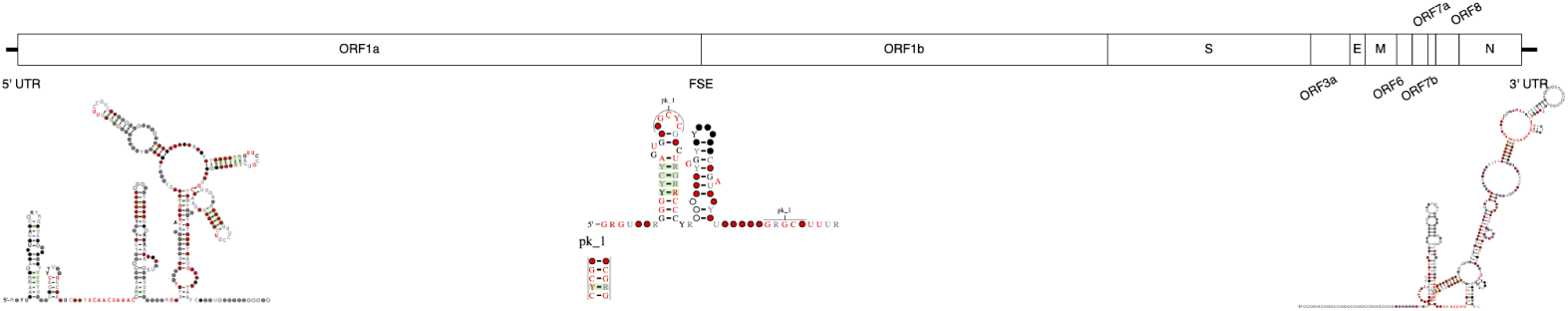
Predicted structures of the 5’ and 3’ UTRs and the frameshifting stimulatory element (FSE) of SARS-CoV-2.

Internal to the SARS-CoV-2 genome and located at the junction between ORF1a and ORF1b, the FSE acts to establish the correct reading frame for ORF1b and all downstream genes by causing an actively translating ribosome to shift back by one nucleotide [15]. Structurally the FSE consists of a seven nucleotide ‘slippery site’, where the ribosome slippage occurs, followed by a five nucleotide spacer, and a pseudoknot that may act as a barrier to the proceeding ribosome, causing slippage within the slippery site approximately 15-30% of the time [5, 15, 27].

### Genomic structure prediction

Since the pandemic onset, work to characterize the secondary structure of SARS-CoV-2 both on a biochemical and a computational basis has been steady and continuous.

Focusing on those computational approaches, predictions of SARS-CoV-2’s secondary structure were released by the Moss laboratory in early 2021, which included the details of their ScanFold-based computational pipeline identifying eight highly likely structures within the genome [2]. At nearly the same time the Pyle laboratory, using the SuperFold RNA secondary structure prediction utility, identified nearly 61% of the viral genome as being base paired and thus exhibiting secondary structure [57]. The Das laboratory, using a combination of CONTRAfold and RNAz identified 44 loci of predicted structure including components of the prototypical 5’ untranslated region (UTR), frameshifting stimulatory element (FSE) and 3’ UTR [12, 49]. The Mathews research group has published their findings using a novel iteration of their TurboFold algorithm, LinearTurboFold, and identified 50 structural elements, 26 of which were novel at the time of publication [32].

Contemporaneously, we employed a combination of RNAz, LocARNA, and CaCoFold to identify secondary structure on the basis of conserved sequence and thermodynamic properties; we identified 40 regions of the genome as very favourable to the formation of secondary structure [18, 50, 69, 75]. These three utilities were chosen for their strengths in identifying likely secondary structure due to thermodynamic properties, both in pseudknot-free and pseudoknotted structures. Further, all three operate on multiple sequence alignment input, allowing us to leverage homology between coronaviruses to identify conserved regions of secondary structure. In that work we additionally compared our findings with a subset of previously published biochemical and computational findings regarding SARS-CoV-2 genome structure. We found that the previously characterized 5’ and 3’ UTRs but also intragenic regions of the genome were likely to generate structure. Further, we found that biochemically based approaches tended to obtain a more similar prediction to one another, while computational approaches varied depending on the algorithmic methods employed.

The goal of the research described here is twofold: to canvas major SARS-CoV-2 variants for synonymous mutations, and to characterize the impact on RNA secondary structure of both synonymous and nonsynonymous mutations on regions we previously predicted as likely to form organized structure. Our hypothesis is that if RNA secondary structure is important to viability of the SARS-CoV-2 virus, synonymous mutations will not lead to major impact on predicted genomic secondary structure. Conversely, if these predicted structures are not important to the viral life cycle and transmissibility, synonymous mutations will be as likely to cause severe changes as they are to cause negligible changes, indicating viral viability is preserved even when predicted structures are not. By measuring mutational rates, both synonymous and non-synonymous, and assessing their impact on predicted structure, we can test this hypothesis.

## 1 Materials and Methods

### 1.1 Data

The reference sequence for SARS-CoV-2, NC 045512.2, was obtained through NCBI’s Nucleotide database [42]. All other sequences were obtained through GISAID’s EpiCoV database [16]. All sequences analyzed are available at https://zenodo.org/doi/10.5281/zenodo.10070695.

### 1.2 Structure prediction

We previously employed a computational pipeline to identify 40 regions likely to form secondary structure within the SARS-CoV-2 genome [18, 50, 69, 75]. Figure 3 summarizes the workflow performed by our pipeline. In summary, thirteen coronavirus genomic sequences including SARS-CoV-2 were aligned to identify regions of the SARS-CoV-2 genome conserved in two or more additional coronavirus genomes. These conserved regions were then analyzed using RNAz, LocARNA, and CaCoFold to identify thermodynamically or stochastic context-free grammatically (SCFG) stable secondary structures as well as covariant nucleotide positions within those structures.

**Fig 3.**
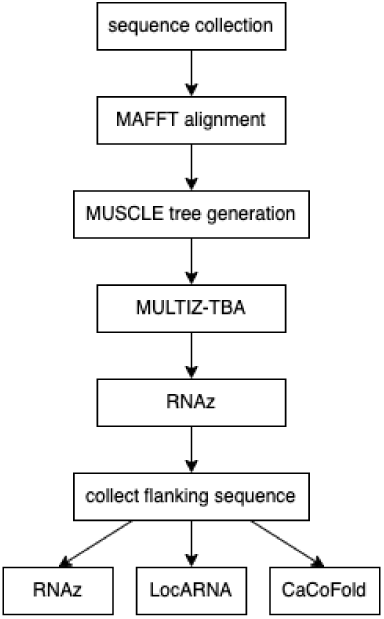
Flowchart describing the methods employed in our previous work.

### 1.3 VoC synonymous mutation identification

For each of the VoCs Alpha, Beta, Gamma, Delta, and Omicron, at least 1000 whole genome samples were collected from GISAID on 14 January 2022. Samples for Omicron BA.2 were collected on 12 March 2022. At the time of collection, all available high coverage, complete Omicron sequences were collected for analysis. For all samples, the following criteria were imposed: the sample was flagged by GISAID as being a member of a given VoC; samples were marked as ‘high coverage’ with low coverage samples further excluded; samples must cover the entire genome of SARS-CoV-2; collection data (including date of collection, submission, and geographic location) was complete for each sample. The rationale for these restrictions was to gather only complete, high quality viral sequences for downstream analysis. Filtered results were sorted on descending sequence length to shuffle geographic and temporal origins of samples. Table 1 describes the number of samples collected for each VoC.

**Table 1.** Number of complete, high coverage viral genomes collected per Variant of Concern.

| VoC | Number of samples |
| --- | --- |
| Alpha | 1006 |
| Beta | 1009 |
| Gamma | 1012 |
| Delta | 1007 |
| Omicron | 5614 |
| Omicron BA.2 | 1023 |

For each VoC collection, multiple sequence alignment was performed using MAFFT with default parameters [26]. We developed Python utilities to parse the resulting multiple sequence alignment at each nucleotide position as compared to the reference genome for SARS-CoV-2, NC 045512.2. This was done to identify all mutations relative to the reference genome for all viral sequences collected. For each position, each observed nucleotide variant was assessed for its frequency within the collected sequences for that VoC and variants observed in at least 75% of the sequences for each VoC were retained; this cutoff was chosen to facilitate comparison with publicly available data offered by outbreak.info, a website specializing in curating SARS-CoV-2 variation data [43]. Outbreak.info reports protein-altering mutations present in at least 75% of the sequences they analyze for each VoC/VoI. Next, a Python tool was developed to characterize each retained nucleotide level variant in terms of its impact on the SARS-CoV-2 proteome to identify synonymous and non-synonymous mutations. For each variant, we determined which codon and codon position that nucleotide variation occupied, identified the reference codon-specified amino acid present in the wild-type protein, and then determined the impact that the detected variation had on that codon. If a nucleotide variation did not alter the amino acid specified at that position, the variation was described as synonymous. Conversely, if a nucleotide variation did alter the amino acid specified at that position, the variation was classified as non-synonymous. A companion Python tool was written to convert publicized protein level variants to nucleotide level mutations; for each amino acid substitution, we determined which nucleotide must have mutated to cause a change from the reference to the mutant amino acid. Figure 4 characterizes the workflow of assessing each mutation for its characteristics including frequency and impact on protein sequence. In summary, mutations present in at least 75% of the samples analyzed were characterized in terms of their synonymous versus non-synonymous status, as well as their presence in the public data repository outbreak.info.

**Fig 4.**
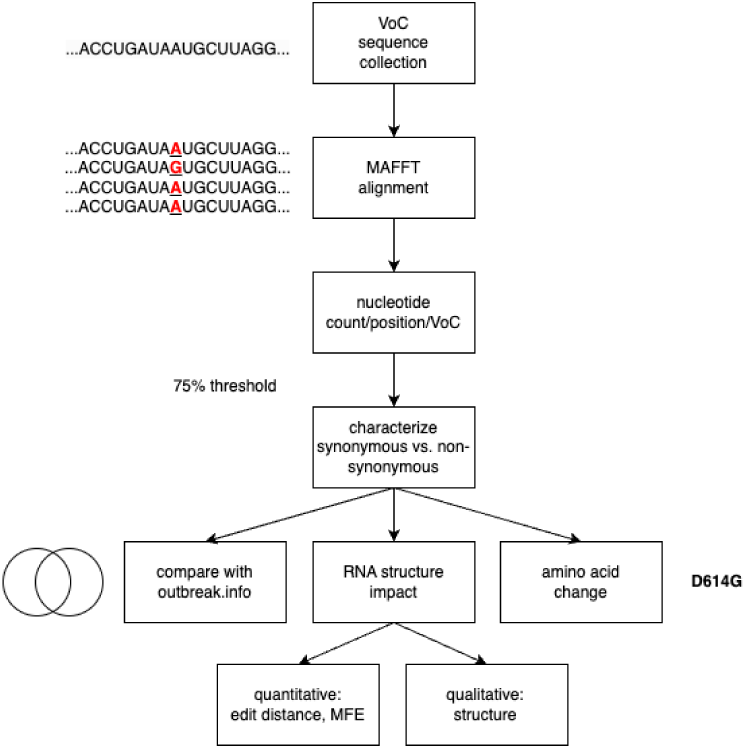
Flowchart describing the methods employed to analyze each Variant of Concern.

### 1.4 Structural impact

All mutations present in at least 75% of the samples analyzed were then assessed for their impact on potential RNA secondary structure formed by the SARS-CoV-2 genome. In our previous research, we identified 40 regions of the viral genome that are most likely to form RNA secondary structures. This identification was based on sequence conservation, predicted thermodynamics, and nucleotide covariation. We then examined the mutations that occurred within these regions, as identified by this work [75]. Those mutations that were present within these 40 regions were then substituted into the reference genome sequence and the mutated subgenomic sequence was checked for structural consequences using RNAfold [36]. We elected to use RNAfold as it is a fast, inexpensive thermodynamic-based method that could quickly assess for any changes in predicted structure. The impact of each mutation on structure was assessed quantitatively by determining the MFE of the reference and mutant structures, and by measuring the Levenshtein string edit distance between the dot-bracket representations of the reference and mutant structures. Levenshtein string edit distance is a measure of the similarity of two sequences of symbols, and quantifies the number of changes needed to convert one sequence to the other. We implemented an algorithm to calculate Levenshtein distance in Python and quantified distances between wild-type and mutant dot bracket structural representations. Impact of mutated positions were assessed qualitatively through visualization using the secondary structure visualization utility VARNA [9]. Fig 4 provides a graphical overview of the methods outlined in this and the previous subsection.

### 1.5 Comparison with public data resources

Outbreak.info is a regularly updated, global repository of data on SARS-CoV-2 variants in circulation currently and historically [43]. Among the data resources available at their website are pages dedicated to describing VoCs/VoIs in terms of their protein-altering (non-synonymous) mutations. Non-synonymous mutations present in at least 75% of the sequences analyzed by outbreak.info are included as ’Characteristic Mutations’ for a given variant. As mutations to UTRs and intergenic regions are not protein-altering, these mutations, along with synonymous mutations, are not included in this report. We compared mutations we identified at the same prevalence of 75% with those catalogued by outbreak.info to measure overlap in identification rates for non-synonymous mutations.

## Results

### 1.6 Characterizations of VoCs

For each of the six VoCs considered, we describe the synonymous and non-synonymous mutations in both coding and non-coding regions detected in at least 75% of sequences. We report mutations by their nucleotide level change and position, and in the case of coding sequence mutations, in terms of their protein level change and position. The following sections detail our findings, organized by VoC. Supplementary table 1 summarizes our findings organized by mutation. We then compare our findings to a publicly available resource, and conclude by describing the impact of identified mutations on predicted secondary structure.

#### 1.6.1 Alpha

Seven synonymous and one non-coding variant were detected in this analysis of SARS-CoV-2 Alpha sequences. Additionally, twenty non-synonymous variants were identified. The overlap between outbreak.info’s reported non-synonymous mutations for Alpha and all variants identified in this work is described in Fig 5. C241U, a mutation in the 5’ UTR was identified as were the following synonymous mutations: in ORF1a, C913U (S216S), C3037U (F924F), C5986U (F1907F); in ORF1b, C14676U (P403P), C15279U (H604H), U16176C (T903T); and in N, G28882A (R203R).

**Fig 5.**
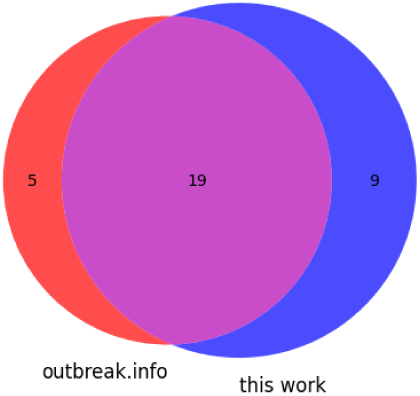
Venn diagram showing the distribution of mutations found in the Alpha variant according to Outbreak.info and these data.

#### 1.6.2 Beta

Two synonymous and two non-coding variants were detected in Beta sequences. Additionally, 33 non-synonymous variants were identified in this work.

There is slight disagreement as to the position of a deletion in the Spike gene: outbreak.info describes this as a deletion of amino acids 241 through 243, which would correspond to nucleotides 22282-22290, while in this work focused on nucleotide level mutation characterization, this deletion spans 22281 through 22289; both nucleotide deletions lead to the same amino acid sequence. We additionally identified a deletion in more than 75% of analyzed samples spanning nucleotides 11288 through 11296, or deletion of Spike amino acids 3675-3677 (residues SGF). The overlap in identification between outbreak.info and this work is presented in Fig 6. Note that while adjacent deleted nucleotides present in more than 75% of analyzed genomes may represent a contiguous deletion, this cannot be confirmed and so the Venn diagram depicted in Figure 6 treats each nucleotide deletion as a separate mutation. G174U and C241U were identified in the 5’ UTR, while C3037U (F924F of ORF1a) and C28253U (F120F of ORF8) were additionally identified.

**Fig 6.**
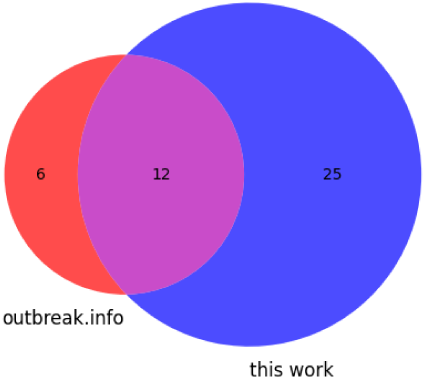
Venn diagram showing the distribution of mutations found in the Beta variant according to Outbreak.info and these data.

#### 1.6.3 Gamma

In the Gamma VoC of SARS-CoV-2, seven synonymous and five non-coding variants were identified, along with 17 non-synonymous mutations. 5’ UTR mutation C241U was again identified as it had been in VoCs Alpha and Beta, as was the insertion of a nucleotide (either A or C) at position 28263 within the intergenic region between ORF8 and N. The synonymous mutations U733C (D156D), C2749U (D828D), C3037U (F924F), A6319G (P2018P), A6613G (V2116V), C12778U (Y4171Y) of ORF1a and C14408U (D131D) of ORF1b were identified. Two non-synonymous mutations not described in outbreak.info were found: ORF1a C3828U (L1191F) was found in 99.8% of samples analyzed, and ORF8 G28167A (E92K) was identified in 96.9% of samples analyzed. The overlap between mutations identified here and by outbreak.info is depicted in Fig 7. Only 35% of the mutations identified were identified by both this work and outbreak.info, with the majority of new mutations being described in this manuscript.

**Fig 7.**
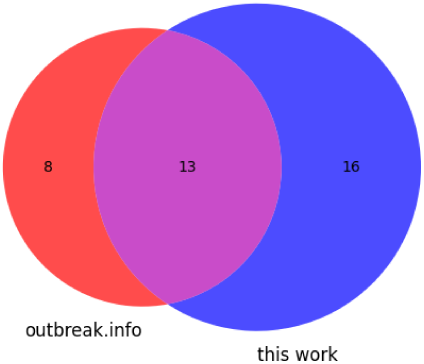
Venn diagram showing the distribution of mutations found in the Gamma variant according to Outbreak.info and these data.

#### 1.6.4 Delta

One synonymous and three non-coding variants were identified in the Delta variant of SARS-CoV-2. G174U and C241U within the 5’ UTR were identified and G29742U within the 3’ UTR, along with C3037U (F924F) within ORF1a. An additional 16 non-synonymous mutations were identified in the Delta samples analyzed. The breakdown of mutations identified by this work and outbreak.info is depicted in Fig 8.

**Fig 8.**
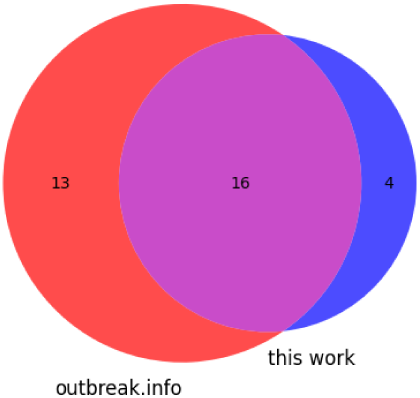
Venn diagram showing the distribution of mutations found in the Delta variant according to Outbreak.info and these data.

Nearly half, 48.5%, of all mutations were identified by both studies, with the majority of the singly identified mutations being found by outbreak.info.

#### 1.6.5 Omicron

At the time of collection, all available full length, high coverage Omicron genomes (5,614) were analyzed. In greater than 75% of samples analyzed, deletions of the 5’ UTR from nucleotide 1 through 28 were observed, with decreasing prevalence ranging from 97.8% deletion (nucleotide position 1) to 75.0% deletion (nucleotide position 28). Similarly, deletions of the 3’ UTR from 29903 (the terminal nucleotide in the genome) up to 29838 were observed, with highest levels of deletion observed in the terminal nucleotide 29903 (96.8% deletion) tapering to 29838 (75.2% deletion).

Additionally the 5’ UTR mutation C241U was observed in Omicron. The synonymous mutations include ORF1a C3037U (F924F), S C25000U (D1146D), ORF3a C25584U (T64T), ORF6 A27259C (R20R) and ORF7b C27807U (L18L). 46 non-synonymous mutations were identified. The overlap between all mutations characterized in this work and mutations reported by outbreak.info is shown in Fig 9. A large proportion of the mutations identified exclusively in this work consist of sequential 5’ and 3’ UTR deletions not reported by outbreak.info.

**Fig 9.**
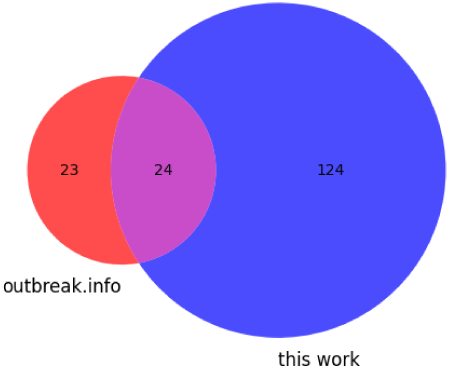
Venn diagram showing the distribution of mutations found in the Omicron variant according to Outbreak.info and these data.

#### 1.6.6 Omicron.BA2

Two non-coding variants and 13 synonymous mutations were identified in analysis of SARS-CoV-2 Omicron.BA2. Fifty-one non-synonymous mutations were identified.

C241U within the 5’ UTR and A28271U within the ORF8/N intergenic boundary were identified, along with the synonymous mutations C3037U (F924F), C4321U (A1352A), A9424G (V3053V), C10198U (D3311D), G10447A (R3394R) and C12880U (I4205I) within ORF1a, C15714U (L749L), A20055G (E2196E) within ORF1b, S mutation C25000U (D1146D), ORF3a mutation C25584U (T64T), M mutation C26858U (F112F), mutation A27259C (R20R) within ORF6 and C27807U (L18L) within ORF7b. Recall that this work treats nucleotide deletions as individual mutations rather than a single contiguous deletion due to the different rates of prevalence of each individual deletion. The overlap between variation identified here and described in outbreak.info is reported in Fig 10. This variant represents the lowest level of overlap in identification, only 20%, with the mutations reported by either individual data source exceeding those reported in common.

**Fig 10.**
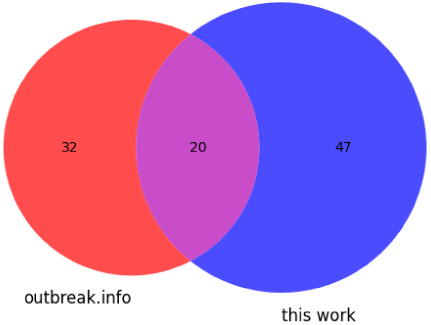
Venn diagram showing the distribution of mutations found in the BA2 variant according to Outbreak.info and these data.

Overall, two mutations were identified at extremely high prevalence in all six VoCs considered, C241U and C3037U. There is a high level of identified synonymous mutations between VoC Omicron and Omicron.BA2, which is understandable as BA2 is derived from its parent Omicron strain. However, the long 5’ and 3’ UTR deletions observed in Omicron samples were not observed in Omicron.BA2 samples.

### 1.7 Comparison with public data resources

In Figure 11, we summarize mutations identified in more than 75% of analyzed samples with those reported coding, non-synonymous mutations according to Outbreak.info, also at a 75% prevalence. For each of the six VoCs, we present the mutation data as ’caterpillar plots’, with our findings on the top half, and Outbreak.info on the bottom half, with the genomic organization of SARS-CoV-2 centred in each plot. For data described in this work, we report deletions as single nucleotide mutations, as our data collection approach typically reports different prevalences for adjacent deleted nucleotides; this is in contrast to the Outbreak.info data which reports deletions as multinucleotide events. In all VoCs and for both data collections, mutations appear more frequently in the Spike, other structural, and 3’ non-structural proteins.

**Fig 11.**
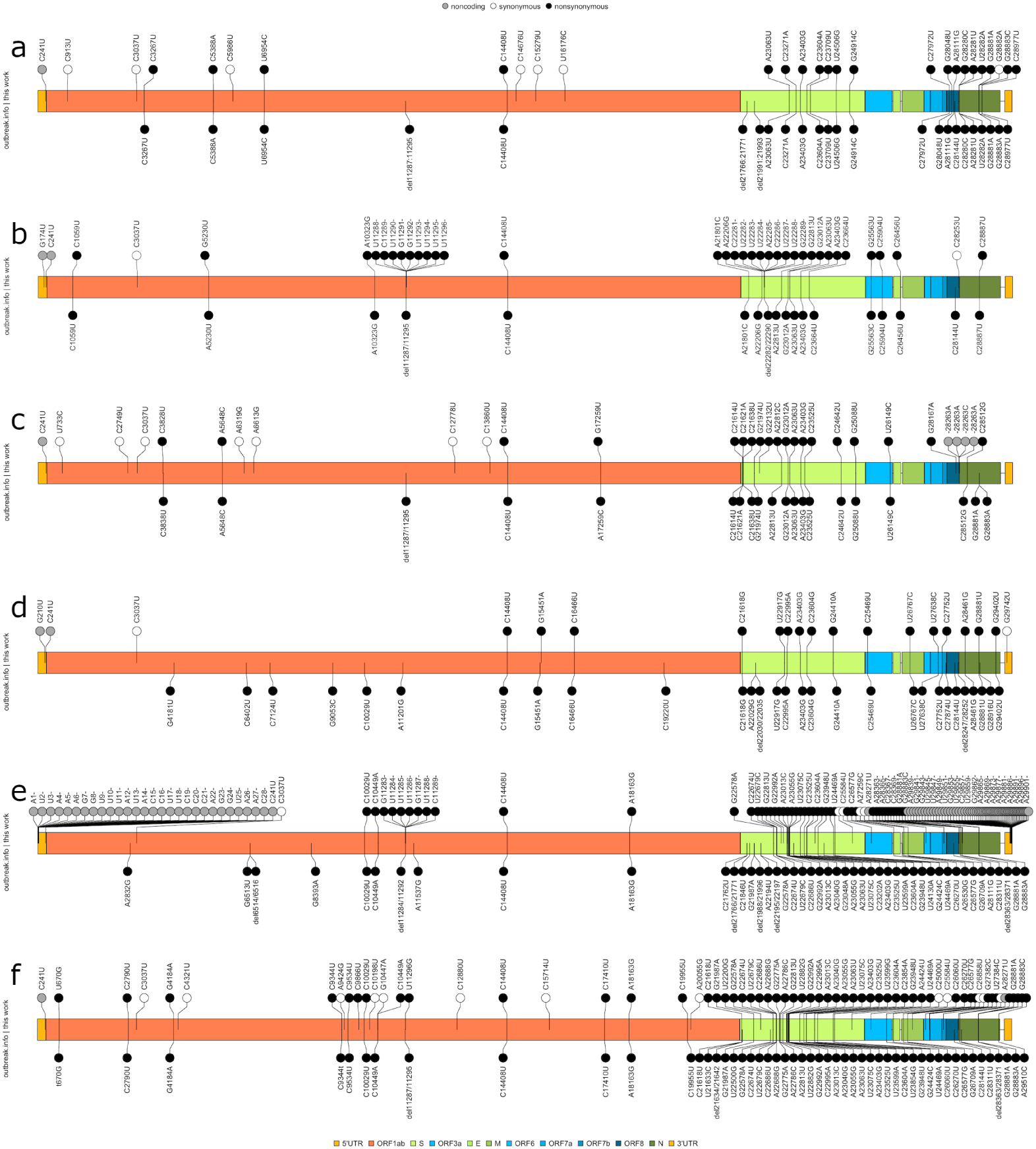
Caterpillar plots comparing non-coding, synonymous, and non-synonymous mutations identified by this work (top row) against those non-synonymous mutations catalogued by Outbreak.info (bottom row). Panels A-F: Alpha, Beta, Gamma, Delta, Omicron and Omicron BA2 VoC mutation profiles.

Qualitatively, it is notable that our data set includes an excess of non-synonymous mutations relative to synonymous mutations; the relative dearth of non-coding mutations is less consequential due to the limited non-coding regions within the viral genome (265 nucleotides for the 5’ UTR, 229 nucleotides for the 3’ UTR, and a total of 284 intergenic non-coding nucleotides, with most intergenic gaps being less than 20 nucleotides in length). When considering only the non-synonymous mutations reported by Outbreak.info, there is not an appreciable but not complete overlap in identified mutations with our data, as is described in the preceding Venn diagrams for each VoC. In some cases, mutations are uniquely reported by Outbreak.info, or by our data. This may be due to the relatively smaller sample size in our data relative to that employed by Outbreak.info in conjunction with the 75% prevalence threshold imposed.

### 1.8 Impact of mutation on predicted structure

For each VoC characteristic mutation identified, the impact of the mutation on structures predicted by our previous work was assessed [75]. An initial observation is that the majority of the mutations identified do not fall within regions we previously predicted as capable of forming secondary structure. In VoC Alpha, 5/28 fall within predicted structure forming regions; for VoC Beta, 6/37; for VoC Gamma, 6/31; for VoC Delta, 4/20; for VoC Omicron, 32/148 and for VoC Omicron.BA2, 12/67. However, the regions predicted to form secondary structure cover 20.9% of the total length of the SARS-CoV-2 genome; a Chi-squared test finds these values do not differ significantly from the coverage value. (Note that for VoC Omicron, the high number of observed mutations is due in large part to the 5’ and 3’ deletions observed in this VoC only: each deleted nucleotide is treated as an individual mutation.) Of the 53 mutations identified among all six VoCs that fell within regions predicted to form secondary structure, 45 of them resulted in quantitative differences in the RNA secondary structure MFE and string edit distance in the mutant sequence relative to the reference sequence. Among those that changed the RNA secondary structure, the mean Levenshtein string edit distance was 26.667, and the mean change in MFE was -0.635. 7 of the 45 structure changing mutations resulted in a more thermodynamically stable structure; that is, the MFE was a negative score of greater magnitude in the mutant than wild-type sequence. Notably the two mutations identified at extremely high prevalence in all six VoCs, C241U and C3037U, did not lead to a quantitative difference in the RNA secondary structure, in terms of both MFE and string edit distance, although they were present in regions predicted to be capable of forming secondary structure. Synonymous and non-coding mutations that change the structure do not have a significantly different change in MFE or Levenshtein distance from non-synonymous mutations.

Supplementary table 1 provides greater detail regarding the previous summary findings.

In terms of the qualitative consequences to structure, the impact of observed mutations range from non-existent: eight of the mutations did not alter structure in any way, to minimal: U26149C, a non-synonymous mutation in ORF3a, reduces a short stem structure by a single base pair, to consequential: GAU27382-27384CUC, a trio of adjacent mutations in ORF6 that leads to an amino acid substitution, causes a longer range rearrangement of a branched stem loop structure (Figs 12 and 13). Other rearrangements include formations of alternate stem-loop structures in existing branched stem loop structures, such as the synonymous mutation C27807U in ORF7b, or C25584U in ORF3a, that decomposes a larger branched stem loop into three adjacent bulged stem-loop structures. The del:1-28 deletion observed in greater than 75% of analyzed Omicron genomes leads to the ablation of SL1 of the predicted 5’ UTR structure. More complex alterations to predicted structure are positively associated with longer Levenshtein distance between the structures and increased absolute change in MFE.

**Fig 12.**
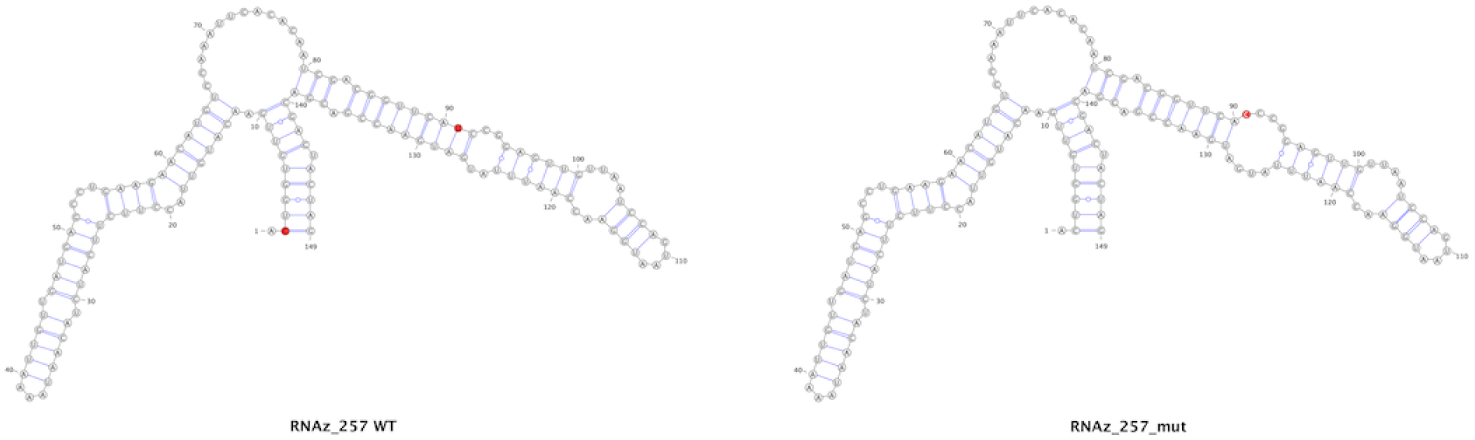
Wildtype (left) and mutated (right) versions of a structure predicted to occur within ORF3a. The red nucleotides indicate the mutated position.

**Fig 13.**
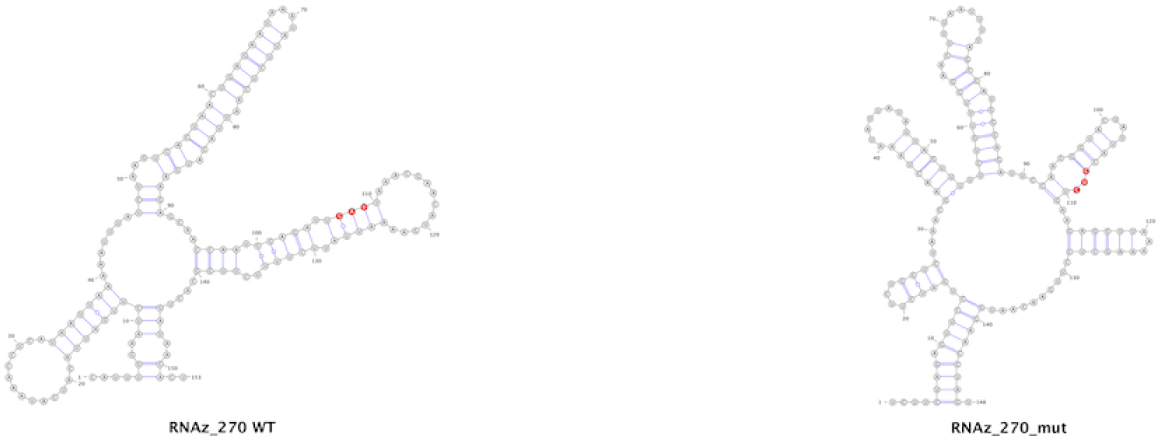
Wildtype (left) and mutated (right) versions of a structure predicted to occur within ORF3a. The red nucleotides indicate the mutated position. Note that this structure prediction is for three adjacent mutated nucleotides.

To summarize, we set out to identify all mutations, including non-coding and synonymous mutations, within the six VoCs of the SARS-CoV-2 genome, with the hypothesis that regions of predicted RNA secondary structure would be especially sensitive to mutations that interfere with that structure formation. We identified many but not all non-synonymous mutations in SARS-CoV-2 VoCs in common with a publicly available data resource. Additionally, we identified mutations to non-coding regions of the genome, as well as synonymous mutations that leave encoded protein sequences unaffected. We identify different patterns of mutation in each VoC, but a limited number of mutations appear to be common to all six analyzed VoCs. Interestingly, these common mutations appear to not impact predicted genome secondary structure configurations. When considering all mutations in terms of their impact to genomic structure, most mutations identified to not impact regions we previously predicted to form secondary structure, and among those that do, the impact to structure is predicted to be modest with few exceptions. Most structure-impacting mutations are mildly destabilizing, causing a predicted increase to MFE. This supports our hypothesis that these regions of predicted structure are biologically relevant to the life cycle of this virus; if these regions of predicted structure were not biologically relevant, we should see a greater frequency of mutations that cause a more serious derangement of predicted structure. Additionally, as we observe far fewer synonymous than non-synonymous mutations, this suggests that RNA sequence is especially important to correct structure formation, perhaps as important in some instances than impact to protein sequence.

## Discussion

As COVID-19 has continued to generate an ongoing global health burden, a number of major VoCs have arisen and dominated the infectious landscape in turn. Largely discussed in terms of their protein-altering mutations, a focus which is not unjustified given the involvement of surface protein structure in the function of monoclonal antibody-based therapeutics and vaccines, the characterization of synonymous mutations, whose impact would be largely restricted to the RNA secondary structure of the viral genome, have only been explored in passing. Additionally, the impact of non-synonymous mutations have been investigated primarily in terms of their protein modifying effects and rarely in terms of their consequences to the genomic secondary structure. We sought to examine the effect of non-coding, synonymous, and non-synonymous mutations on the RNA secondary structure of the SARS-CoV-2 genome within the framework of six prominent VoCs. Underlying this investigation was the question as to whether mutations, regardless of their protein-altering potential, could impact the integrity of regions we’ve previously predicted as capable of generating RNA secondary structure. If identified mutations routinely altered those predicted structures, it would argue against the importance of those structures for the viability of SARS-CoV-2. Conversely, the absence or infrequent occurrence of structure altering mutations would leave the possibility intact that those structures are indeed significant.

In all variants examined, the number of non-synonymous mutations identified among at least 75% of genomes analyzed consistently exceeds the number of non-coding and synonymous mutations. The relative dearth of non-coding mutations is in no small part due to the very restricted number of non-coding nucleotides in the SARS-CoV-2 genome, but is also likely due to the known functional structures in the 5’ and 3’ UTR: mutations in these regions are very likely to impact structure and so impair those functions. Nonsense mutations (protein altering mutations that change an amino acid to a stop codon) are nearly absent, with a single exception of C27972U (ORF8, Q27Ter) in VoC Alpha; this mutation does not land in a predicted structure-forming region. As we are only considering mutations present in at least 75% of analyzed genomes, this work does not lead to a meaningful calculation of the ratio of non-synonymous to synonymous mutations (dN/dS) for SARS-CoV-2 in its entirety or for individual genes, but the preponderance of non-synonymous mutations is certainly suggestive of a ratio greater than 1, implying overall positive selection on the genome. Had we observed a surplus of synonymous mutations, the result could be qualitatively described as negative or purifying selection.

There has been some limited work on non-protein affecting mutations. In a 2020 paper, Kannan and colleagues observe the high frequency of C241U and speculate that it may alter the structure of SL5 in the 5’ UTR or impact protein binding in that region [25]. A paper published by Ryder *et al.* in 2021 also observes the relatively high frequency of C241U and infers the possibility that it is induced by host RNA editing [52]. In a series of computational analyses, Chaudhari and colleagues generate potential three dimensional structures for the wild type and C241U version of the 5’ UTR and find both an RNA structural change and alteration in predicted docking with the transcription factor MADP1 [8]. Similarly the C3037U synonymous mutation has also previously been discussed. In 2020 van Dorp reported that C3037U is homoplasic and in linkage disequilibrium with A23403G (S, G614D) and in the same year it was reported that C3037U disrupts a potential binding site for the miRNA miR-197-5p, a miRNA previously reported to be elevated in influenza H7N9 patients [21, 45, 62]. We find that C241U imposes no quantitative or qualitative change to predicted structure or MFE. In terms of C3037U, we similarly find no impact to predicted secondary structure in this region of the genome. Presumably the non-impact to structure, in conjunction with the non-coding and synonymous natures of C241U and C3037U respectively, allow these mutations to be truly invisible to selective pressure themselves, but as in the case of C3037U, may be passenger mutations following selection on linked, selected mutations (A23403G in that case).

Most recently, a 2023 paper by Bai *et al.* describes a limited characterization of the influence of synonymous mutations on potential RNA structure; contrary to our findings, that work reports a dN/dS ratio of less than 1, indicating purifying selection, and observes a roughly even distribution of synonymous mutations throughout the genome [3]. In this work, the authors analyze 6483 viral genomes in terms of their synonymous and non-synonymous mutation occurences as compared to the reference genome for the purposes of assessing codon usage changes over time. To this end they collect data on all mutations, not only those that exceed a threshold as is done in our work. Further, their RNA secondary structure prediction data is derived from a single DMS-MaP-seq experiment, rather than from a structural conservation-based approach as undertaken in our previous work [40]. Bai’s observations lead the authors to conclude that synonymous mutations are unlikely to impinge upon RNA secondary structure function, as they do not observe ’hot’ or ’cold’ spots for synonymous mutations. We argue that there is far greater potential for secondary structure across the entire genomic span including coding regions, and that the relative dearth of synonymous mutations is suggestive of the importance of those structures. We suggest that the fundamental differences in findings are due in large part to the differences in both mutation calculation and structure prediction, and that as a result, these findings are not appropriate for direct comparison with the work presented here.

In this work long terminal deletions of both the 5’ and 3’ UTRs are observed in the majority of Omicron samples, approaching 98% prevalence at some nucleotide positions. While it would be reasonable to exclude these samples as incomplete, at the time of collection all Omicron samples listed as full sequence, high coverage were included, 5614 full genomes. These samples were obtained from all populated continents and a wide variety of countries. It seems improbable that nearly all of these represent miscategorized partial sequence genomes, but that possibility may not be excluded. De Maio and colleagues recommend masking the first 55 and last 100 nucleotides of SARS-CoV-2 for subsequent analysis, recognizing the possibility of low sequence coverage generating aberrant results [10]. In the case that these do represent true complete sequences, there are consequences to predicted structure. For the 5’ end of the genome, twenty eight nucleotides are absent in more than 75% of the Omicron samples analyzed, leading to the removal of 5’ UTR SL1. This absence is notable, due to SL1’s role in evading translational repression [6]. Loss of this structure would presumably render these del:1-28 Omicron sequences susceptible to Nsp1-mediated repression.

Similarly del:29838-29903 observed in the Omicron sequences would potentially influence the 3’ UTR structure, although this deletion is downstream from any previously identified structure; our previous work does not characterize this region as likely of forming secondary structure, either. We cannot rule out the possibility that these are spurious ‘mutations’, the result of sequencing error or degraded sequence due to compromised sample handling, although we have attempted to avoid these possibilities by restricting our samples to high quality, high coverage, full length genomic sequence.

## Conclusion

In this work we present our efforts to characterize the SARS-CoV-2 genome in terms of its high frequency RNA genome structure-altering mutations in six VoCs. We anticipated that if regions we previously predicted to generate RNA secondary structure were important to viral biology, we would be less likely to find mutations that negatively affect the integrity of those predicted structures. Conversely, if these structures were not relevant, we would see more synonymous mutations that maintain a wild type protein sequence, effectively being an invisible mutation with regards to selection. We find a surplus of non-synonymous, protein-altering mutations relative to synonymous mutations; and we infer that selective pressure is likely positive.

Predictions of the effects of identified mutations that do affect structure at all typically have a modest or even negligible effect, suggesting the importance of RNA secondary structure to the life cycle of the virus. Regions of predicted structure within coding regions of the viral genome show a similar lack of structure-altering mutations, implying that genome structure is as important in genic regions as in untranslated regions. These findings support the notion that these structural regions are likely important to viral biology, to such a degree that even synonymous mutations are at risk of impairing viral fitness. Additionally, we see few mutations in non-coding regions, and those we do identify similarly have non-existent or negligible impact to UTR structure, with the key exception of detected Omicron terminal deletions; it remains to be determined if these are of biological or technical origin. A better understanding of this pandemic causing virus in terms of its genomic structure will lead to improved, accelerated assessment of future RNA virus outbreaks.

## Supporting information

**S1 File. Summary of variant positions identified within our previously determined regions of structure.** WT MFE: wild type (reference) structure minimum free energy. Mut MFE: mutant/variant structure minimum free energy. L dist: Levenshtein distance.

## Acknowledgments

This work was supported in part by a grant from Microsoft Azure AI for Health.

## References

1. Andersen K, Rambaut A, Lipkin WI, Holmes E, Garry R. The proximal origin of SARS-CoV-2. Nature Medicine 2020 Apr;26(4):450–2.

2. Andrews R, O’Leary C, Tompkins V, Peterson J, Haniff H, Williams C, Disney M, Moss W. A map of the SARS-CoV-2 RNA structurome. Nucleic Acids Research Genomics and Bioinformatics 2021 Jun 1;3(2):lqab043.

3. Bai H, Ata H, Sun Q, Rahman SU, Tao S. Natural selection pressure exerted on ”silent” mutations during the evolution of SARS-CoV-2: evidence from codon usage and RNA structure. Virus Research 2023 Jan 2;323:198966.

4. Bal A, Destras G, Gaymard A, Stefic K, Marlet J, Eymieux S, Regue H, Semanas Q, d’Aubarede C, Billaud G, Laurent F, Gonzalez C, Mekki Y, Valette M, Bouscambert M, Gaudy-Graffin C, Lina B, Morfin F, Josset L, COVID-Diagnosis HCL Study Group. Two-step strategy for the identification of SARS-CoV-2 variant of concern 202012/01 and other variants with spike deletion H69-V70, France, August to December 2020. Eurosurveillance 2021 Jan 21;26(3):2100008.

5. Brierley I, Pennell S, Gilbert R. Viral RNA pseudoknots: versatile motifs in gene expression and replication. Nature Reviews Microbiology 2007 Aug;5(8):598–610.

6. Bujanic L, Shevchuk O, von Kugelgen N, Kalinina A, Ludwik K, Koppstein D, Zerna N, Sickmann A, Chekulaeva M. The key features of SARS-CoV-2 leader and NSP1 required for viral escape of NSP1-mediated repression. em RNA 2022 May 1;28(5):766–79.

7. Chan KK, Tan TJC, Narayanan KK, Procko E. An engineered decoy receptor for SARS-CoV-2 broadly binds protein S sequence variants. Science Advances 2021 Feb 17;7(8):eabf1738.

8. Chaudhari A, Chaudhari M, Mahera S, Saiyed Z, Nathani NM, Shukla S, Patel D, Patel C, Joshi M, Joshi CG. In-silico analysis reveals lower transcription efficiency of C241T variant of SARS-CoV-2 with host replication factors MADP1 and hnRNP-1. Informatics in Medicine Unlocked 2021 Jan 1;25:100670.

9. Darty K, Denise A, Ponty Y. VARNA: interactive drawing and editing of the RNA secondary structure. Bioinformatics 2009 Aug 8;25(15):1974.

10. Issues with SARS-CoV-2 sequencing data. https://virological.org/t/issues-with-sars-cov-2-sequencing-data/473 Accessed 15 July 2023.

11. Di Giacomo S, Mercatelli D, Rakhimov A, Giorgi FM. Preliminary report on severe acute respiratory syndrome coronavirus 2 (SARS-CoV-2) Spike mutation T478K. Journal of Medical Virology 2021 Sep;93(9):5638–43.

12. Do CB, Woods DA, Batzoglou S. Contrafold: RNA secondary structure prediction without physics-based models. Bioinformatics 2006 Jul 15;22(14):e90–8.

13. Faria NR, Mellan TA, Whittaker C, Claro IM, Candido DDS, Mishra S, Crispim MA, Sales FC, Hawryluk I, McCrone JT, Hulswit RJ. Genomics and epidemiology of the P.1 SARS-CoV-2 lineage in Manaus, Brazil. Science 2021 May 21;372(6544):815–21.

14. Gangavarapu K, Latif AA, Mullen JL, Alkuzweny M, Hufbauer E, Tsueng G, Haag E, Zeller M, Aceves CM, Zaiets K, Cano M, Zhou X, Qian Z, Sattler R, Matteson NL, Levy JI, Lee RTC, Freitas L, Maurer-Stroh S, GISAID Core and Curation Team, Suchard MA, Wu C, Su AI, Andersen KG, Hughes LD. Outbreak.info genomic reports: scalable and dynamic surveillance of SARS-CoV-2 variants and mutations. Nature Methods 2023 Apr;20(4):512–22.

15. Giedroc D, Cornish P. Frameshifting RNA pseudoknots: structure and mechanism. Virus Research 2009 Feb 1;139(2):193–208.

16. GISAID - hCov19 Variants https://www.gisaid.org/hcov19-variants/ Accessed 13 July 2023.

17. Goebel S, Hsue B, Dombrowski T, Masters P. Characterization of the RNA components of a putative molecular switch in the 3’ untranslated region of the murine coronavirus genome. Journal of Virology 2004 Jan 15;78(2):669–82.

18. Gruber A, Findeis S, Washietl S, Hofacker I, Stadler P. RNAZ 2.0: improved noncoding RNA detection. In Biocomputing 2010 pp.69–79.

19. Gu H, Chen Q, Yang G, He L, Fan H, Deng YQ, Wang Y, Teng Y, Zhao Z, Chi Y, Li Y. Adaptation of SARS-CoV-2 in BALB/c mice for testing vaccine efficacy. Science 2020 Sep 25;369(6511):1603-7.

20. Guan BJ, Su YP, Wu HY, Brian D. Genetic evidence of a long range RNA-RNA interaction between the genomic 5’ untranslated region and the nonstructural protein 1 coding region in murine and bovine coronaviruses. Journal of Virology 2012 Apr 15;86(8):4631–43.

21. Hosseini Rad SM A, McLellan AD. Implications of SARS-CoV-2 mutations for genomic RNA structure and host microRNA targeting. International Journal of Molecular Sciences 2020 Jul 7;21(13):4807.

22. Hsue B, Masters PS. A bulged stem-loop structure in the 3’ untranslated region of the genome of the coronavirus mouse hepatitis virus is essential for replication. Journal of Virology 1997 Oct;71(10):7567–78.

23. Hsue B, Hartshorne T, Masters P. Characterization of an essential RNA secondary structure in the 3’ untranslated region of the murine coronavirus genome. Journal of Virology 2000 Aug 1;74(15):6911–21.

24. Hu B, Guo H, Zhou P, Shi ZL. Characteristics of SARS-CoV-2 and covid-19. Nature Reviews Microbiology 2021 Mar;19(3):141–54.

25. Kannan SR, Spratt AN, Quinn TP, Heng X, Lorson CL, Sonnerborg A, Byrareddy SN, Singh K. Infectivity of SARS-CoV-2: there is something more than D614G? Journal of Neuroimmune Pharmacology 2020 Dec;15:574–7.

26. Kato K, Standley D. MAFFT multiple sequence alignment software version 7: improvements in performance and usability. Molecular Biology and Evolution 2013 Jan 16;30(4):772–80.

27. Kelly J, Olson A, Neupane K, Munshi S, San Emeterio J, Pollack L, Woodside M, Dinman J. Structural and functional conservation of the programmed -1 ribosomal frameshift signal of SARS coronavirus 2 (SARS-CoV-2). Journal of Biological Chemistry 2020 Jul 31;295(31):10741–8.

28. Khan A, Zia T, Suleman M, Khan T, Ali SS, Abbasi AA, Mohammad A, Wei DQ. Higher infectivity of the SARS-CoV-2 new variants is associated with K417N/T, E484K, and N501Y mutants: an insight from structural data. Journal of Cell Physiology 2021 Oct;236(10):7045–57.

29. Lee CW, Li L, Geidroc D. The solution structure of coronaviral stem-loop 2 (SL2) reveals a canonical CUYG tetraloop fold. FEBS Letters 2011 Apr 6;585(7):1049–53.

30. Li W, Moore MJ, Vasilieva N, Sui J, Wong SK, Berne MA, Somasundaran M, Sullivan JL, Luzuriaga K, Greenough TC, Choe H, Farzan M. Angiotensin-converting enzyme 2 is a functional receptor for the SARS coronavirus. Nature 2003 Nov 27;426(6965):450-4.

31. Li L, Kang H, Liu P, Makkinje N, Williamson S, Leibowitz J, Geidroc D. Structural lability in stem-loop 1 drives a 5’UTR-3’UTR interaction in coronavirus replication. Journal of Molecular Biology 2008 Mar 28;377(3):790–803.

32. Li S, Zhang H, Zhang L, Liu K, Liu B, Mathews D, Huang L. LinearTurboFold: linear-time global prediction of conserved structures for RNA homologs with applications to SARS-CoV-2. Proceedings of the National Academy of Sciences 2021 Dec 28;118(52):e2116269118.

33. Liu P, Li L, Millership J, Kang H, Leibowitz J, Geidroc D. A U-turn motif-containing stem-loop in the coronavirus 5’ untranslated region plays a functional role in replication. RNA 2007 May 1;13(5):763–80.

34. Liu Y, Liu J, Plante KS, Plante JA, Xie X, Zhang X, Ku Z, An Z, Scharton D, Schindewolf C, Widen SG, Menachery VD, Shi PY, Weaver SC. The N501Y spike substitution enhances SARS-CoV-2 infection and transmission. Nature 2022 Feb 10;602(7896):294-9.

35. Liu C, Ginn HM, Dejnirattisai W, Supasa P, Wang B, Tuekprakhon A, Nutalai R, Zhou D, Mentzer AJ, Zhao Y, Duyvesteyn HM. Reduced neutralization of SARS-CoV-2 B.1.617 by vaccine and convalescent serum. Cell 2021 Aug 5;184(16):4220–36.

36. Lorenz R, Bernhart S, Honer zu Siederdissen C, Tafer H, Flamm C, Stadler P, Hofacker I. ViennaRNA package 2.0. Algorithms for Molecular Biology 2011 Dec;6:1–4.

37. Lubinski B, Fernandes MHV, Frazier L, Tang T, Daniel S, Diel DG, Jaimes JA, Whittaker GR. Functional evaluation of the P681H mutation on the proteolytic activation of the SARS-CoV-2 variant B.1.1.7 (Alpha) spike. iScience 2022 Jan 21;25(1).

38. Madhugiri R, Karl N, Petersen D, Lamkiewicz K, Fricke M, Wend U, Scheuer R. Structural and functional conservation of cis-acting RNA elements in coronavirus 5’-terminal genome regions. Virology 2018 Apr 1;517:44–55.

39. Meng B, Kemp SA, Papa G, Datir R, Ferreira IATM, Marelli S, Harvey WT, Lytras S. Recurrent emergence of SARS-CoV-2 spike deletion H69/V70 and its role in the Alpha variant B.1.1.7. Cell Reports 2021 Jun 29;35(13).

40. Morandi E, Manfredonia I, Simon LM, Anselmi F, van Hemert MJ, Oliviero S, Incarnato D. Genome-scale deconvolution of RNA structure ensembles. Nature Methods 2021 Mar;18(3):249–252.

41. Mustafa Z, Kalbacher H, Burster T. Occurrence of a novel cleavage site for cathepsin G adjacent to the polybasic sequence within the proteolytically sensitive activation loop of the SARS-CoV-2 Omicron variant: the amino acid substitution N679K and P681H of the spike protein. PLoS One 2022 Apr 18;17(4):e0264723.

42. Home - Nucleotide - NCBI https://www.ncbi.nlm.nih.gov/nucleotide/ Accessed 13 July 2023.

43. outbreak.info SARS-CoV-2 data explorer https://outbreak.info/ Accessed 13 July 2023.

44. Peacock TP, Goldhill DH, Zhou J, Baillon L, Frise R, Swann OC, Kugathasan R, Penn R, Brown JC, Sanchez-David RY, Braga L, Kavanagh Williamson M, Hassard JA, Staller E, Hanley B, Osborn M, Giacca M, Davidson AD, Matthews DA, Barclay WS. The furin cleavage site of SARS-CoV-2 spike protein is a key determinant for transmission due to enhanced replication in airway cells. bioRxiv 2020 Sep 30:2020–09..

45. Peng F, Loo JFC, Kong SK, Li B, Gu D. Identification of serum microRNAs as diagnostic biomarkers for influenza H7N9 infection. Virology Reports 2017 Jun 1;7:1–8.

46. Planas D, Veyer D, Baidaliuk A, Staropoli I, Guivel-Benhassine F, Rajah MM, Planchais C, Porrot F, Robillard N, Puech J, Prot M. Reduced sensitivity of SARS-CoV-2 variant Delta to antibody neutralization. Nature 2021 Aug 12;596(7871):276-80.

47. Qu P, Xu K, Faraone JN, Goodarzi N, Zheng YM, Carlin C, Bednash JS, Horowitz JC, Mallampalli RK, Saif LJ, Oltz EM, Jones D, Gumina RJ, Liu SL. Immune evasion, infectivity, and fusogenicity of SARS-CoV-2 BA.2.86 and FLip variants. Cell 2024 Feb 1;187(3): 585–595.e6.

48. Preliminary genomic characterisation of an emergent SARS-CoV-2 lineage in the UK defined by a novel set of spike mutations. https://virological.org/t/preliminary-genomic-characterisation-of-an-emergent-sars-cov-2-lineage-in-the-uk-defined-by-a-novel-set-of-spike-mutations/563 Accessed 12 July 2023.

49. Rangan R, Zheludev I, Havey R, Pham E, Wayment-Steele H, Glenn J, Das R. RNA genome conservation and secondary structure in SARS-CoV-2 and SARS-related diseases: a first look. RNA 2020 Aug 1;26(8):937–59.

50. Rivas, E. RNA structure prediction using positive and negative evolutionary information. PLoS Computational Biology. 2020 Oct 30;16(10):e1008387.

51. Robertson M, Igel H, Baertsch R, Haussler D, Ares M, Scott W. The structure of a rigorously conserved RNA element within the SARS virus genome. PLoS Biology 2005 Jan;3(1):e5.

52. Ryder SP, Morgan BR, Coskun P, Antkowiak K, Massi F. Analysis of emerging variants in structured regions of the SARS-CoV-2 genome. Evolutionary Bioinformatics 2021 May;17:11769343211014167.

53. Saito A, Takashi I, Suzuki R, Maemura T, Nasser H, Uriu K, Kosugi Y, Shirakawa K, Sadamasu K, Kimura I, Ito J, Wu J, Iwatsuki-Horimoto K, Ito M, Yamayoshi S, Loeber S, Tsuda M, Wang L, Ozono S, Butlertanaka EP, Tanaka YL, Shimizu R, Shimizu K, Yoshimatsu K, The Genotype to Phenotype Japan (G2P-Japan) Consortium, Tanaka S, Nakagawa S, Ikeda T, Fukuhara T, Kawaoka Y, Sato K. Enhanced fusogenicity and pathogenicity of SARS-CoV-2 Delta P681R mutation. Nature 2022 Feb 10;602(7896):300–6.

54. Starr TN, Greaney AJ, Hilton SK, Ellis D, Crawford KHD, Dingens AS, Navarro MJ, Bowen MJ, Tortorici MA, Walls AC, King NP. Deep mutational scanning of SARS-CoV-2 receptor binding domain reveals constraints on folding and ACE2 binding. Cell 2020 Sep 3;182(5):1295–310.

55. Starr TN, Greaney AJ, Dingens AS, Bloom JD. Complete map of SARS-CoV-2 RBD mutations that escape the monoclonal antibody LY-CoV555 and its cocktail with LY-CoV016. Cell Reports Medicine 2021 Apr 20;2(4).

56. Tao K, Tzou PL, Nouhin J, Gupta RK, de Oliveira T, Kosakovsky Pond SL, Fera D, Shafer RW. The biological and clinical significance of emerging SARS-CoV-2 variants. Nature Reviews Genetics 2021 Dec;22(12):757–73.

57. de Cesaris Araujo Tavares R, Mahadeshwa G, Wan H, Huston N, Pyle AM. The global and local distribution of RNA structure throughout the SARS-CoV-2 genome. Journal of Virology 2021 Feb 10;95(5):10–128.

58. Tegally H, Wilkonson E, Giovanetti M, Iranzadeh A, Fonseca V, Giandhari J, Doolabh D, Pillay S, San EJ, Msomi N, Mlisana K. Detection of a SARS-CoV-2 variant of concern in South Africa. Nature 2021 Apr 15;592(7854):438-43.

59. Tian D, Sun Y, Xu H, Ye Q. The emergence and epidemic characteristics of the highly mutated SARS-CoV-2 Omicron variant. Journal of Medical Virology 2022 Jun;94(6):2376–83.

60. Tengs T, Jonassen C. Distribution and evolutionary history of the mobile genetic element s2m in coronaviruses. Diseases 2016 Jul 28;4(3):27.

61. Tsang KW, Ho PL, Ooi GC, Yee WK, Wang T, Chan-Yeung M, Lam WK, Seto WH, Yam LY, Cheung TM, Wong PC. A cluster of cases of Severe Acute Respiratory Syndrome in Hong Kong. New England Journal of Medicine 2003 May 15;348(20):1977–85.

62. van Dorp L, Richard D, Tan CCS, Shaw LP, Acman M, Balloux F. No evidence for increased transmissibility from recurrent mutations in SARS-CoV-2. Nature Communications 2020 Nov 25;11(1):5986.

63. Volz E, Mishra S, Chand M, Barrett JC, Johnson R, Geidelberg L, Hinsley WR, Laydon DJ, Dabrera G, O’Toole A, Amato R. Assessing transmissibility of SARS-CoV-2 lineage B.1.1.7 in England. Nature 2021 May 13;593(7858):266–9.

64. Wan Y, Shang J, Graham R, Baric RS, Li F. Receptor recognition by the novel coronavirus from Wuhan: an analysis based on decade-long structural studies of SARS coronavirus. Journal of Virology 2020 Mar 17;94(7):10–128.

65. Weisblum Y, Schmidt F, Zhang F, DaSilva J, Poston D, Lorenzi JCC, Muecksch F, Rutkowska M, Hoffmann HH, Michailidis E, Gaebler C. Escape from neutralizing antibodies by SARS-CoV-2 spike protein variants. eLife 2020 Oct 28;9:e61312.

66. Updated working definitions and primary actions for SARS-CoV-2 variants. https://www.who.int/publications/m/item/updated-working-definitions-and-primary-actions-for-sars-cov-2-variants Accessed 12 July 2023.

67. Weekly epidemiological update on COVID-19 - 4 May 2021. https://www.who.int/publications/m/item/weekly-epidemiological-update-on-covid-19-4-may-2021 Accessed 13 July 2023.

68. Classification of Omicron (B.1.1.529): SARS-CoV-2 Variant of Concern. https://www.who.int/news/item/26-11-2021-classification-of-omicron-(b.1.1.529)-sars-cov-2-variant-of-concern Accessed 13 July 2023.

69. Will S, Joshi T, Hofacker I, Stadler P, Backofen R. LocARNA-P: accurate boundary prediction and improved detection of structural RNAs. RNA 2012 May 1;18(5):900–14.

70. Williams G, Chang RY, Brian D. A phylogenetically conserved hairpin-type 3’ untranslated region pseudoknot functions in coronavirus RNA replication. Journal of Virology 1999 Oct 1;73(10):8349–55.

71. Wu HY, Guan BJ, Su YP, Fan YH, Brian D. Reselection of a genomic upstream open reading frame in mouse hepatitis coronavirus 5’-untranslated region mutants. Journal of Virology 2014 Jan 15;88(2):846–58.

72. Wu F, Zhao S, Yu B, Chen YM, Wang W, Song ZG, Hu Y, Tao ZW, Tian JH, Pei YY, Yuan ML, Zhang YL, Dai FH, Liu Y, Wang QM, Zheng JJ, Xu L, Holmes E, Zhang YZ. A new coronavirus associated with human respiratory disease in China. Nature 2020 Mar;579(7798):265-9.

73. Yang D, Liu P, Giedroc D, Leibowitz J. Mouse hepatitis virus stem-loop 4 functions as a spacer element required to drive subgenomic RNA synthesis. Journal of Virology 2011 Sep 1;85(17):9199–209.

74. Yang D, Liu P, Wudeck E, Giedroc D, Leibowitz J. SHAPE analysis of the RNA secondary structure of the mouse hepatitis virus 5’ untranslated region and N-terminal Nsp1 coding sequences. Virology 2015 Jan 15;475:15–27.

75. Ziesel A, Jabbari H. Unveiling hidden structural patterns in the SARS-CoV-2 genome: Computational insights and comparative analysis. PLOS ONE 2024 19(4):e0298164.

