## Supplemental Table 1 for "Structural impact of synonymous mutations in six SARS-CoV-2 Variants of Concern"

Table 1: Summary of variant positions identified within our previously determined regions of structure. WT MFE: wild type (reference) structure minimum free energy. Mut MFE: mutant/variant structure minimum free energy. L dist: Levenshtein distance.

| VoC | Structure | Nucleotide | Gene/Region | Protein | WT MFE | Mut MFE | Δ MFE | L dist | In outbreak? |
|---|---|---|---|---|---|---|---|---|---|
| omicron | RNAz_0 | del:1-28 | 5'UTR | | -35.1 | -26.6 | 8.5 | 28 | NO |
| beta | RNAz_1 | g174t | 5'UTR | | -50.7 | -47.7 | 3.0 | 6 | NO |
| delta | RNAz_1 | g210t | 5'UTR | | -50.7 | -49.6 | 1.1 | 18 | NO |
| all | RNAz_2 | c241t | 5'UTR | | -47.5 | -47.5 | 0.0 | 0 | NO |
| alpha | RNAz_9 | c913t | ORF1a | S216S | -44.4 | -41.7 | 2.7 | 31 | NO |
| omicron.BA2 | RNAz_28 | c2790t | ORF1a | T842I | -51.7 | -51.7 | 0.0 | 0 | YES |
| all | RNAz_30 | c3037t | ORF1a | F924F | -45.9 | -45.9 | 0.0 | 0 | NO |
| alpha | RNAz_32 | c3267t | ORF1a | T1000I | -45.3 | -43.4 | 1.9 | 19 | YES |
| omicron.BA2 | RNAz_44 | c4321t | ORF1a | A1352A | -36.8 | -34.9 | 1.9 | 28 | NO |
| beta | RNAz_53 | g5230t | ORF1a | K1655N | -34.8 | -31.3 | 3.5 | 12 | YES |
| alpha | RNAz_54 | c5388a | ORF1a | A1708D | -39.2 | -39.2 | 0.0 | 0 | YES |
| gamma | RNAz_58 | a5648c | ORF1 | K1795Q | -41.3 | -41.3 | 0.0 | 0 | YES |
| gamma | RNAz_67 | a6613g | ORF1a | V2116V | -35.5 | -35.5 | 0.0 | 0 | NO |
| omicron.BA2 | RNAz_220 | g21987a | S | G142D | -27.5 | -25.0 | 2.5 | 56 | YES |
| gamma | RNAz_221 | g22132t | S | R190S | -32.7 | -33.1 | -0.4 | 8 | YES |
| delta | RNAz_250 | c25469t | ORF3a | S26L | 028.7 | -28.7 | 0.0 | 0 | YES |
| omicron, omicron.BA2 | RNAz_252 | c25584t | ORF3a | T64T | -31.9 | -32.8 | -0.9 | 30 | NO |
| beta | RNAz_255 | c25904t | ORF3a | S171L | -43.1 | -43.1 | 0.0 | 0 | YES |
| gamma | RNAz_257 | t26149c | ORF3a | S253P | -46.3 | -42.9 | 3.4 | 4 | YES |
| omicron.BA2 | RNAz_257 | c26060t | ORF3a | T223I | -46.3 | -45.5 | 0.8 | 18 | YES |
| beta | RNAz_261 | c26456t | E | P71L | -23.4 | -23.4 | 0.0 | 0 | YES |
| omicron.BA2 | RNAz_262 | c26577g | M | Q19E | -41.7 | -43.4 | -1.7 | 16 | YES |
| omicron.BA2 | RNAz_270 | gat27382ctc | ORF6 | D61L | -20.5 | -34.3 | -13.8 | 64 | YES |
| omicron, omicron.BA2 | RNAz_275 | c27807t | ORF7b | L18L | -27.2 | -26.3 | 0.9 | 11 | NO |
